# Dissociating physiological ripples and epileptiform discharges with vision transformers

**DOI:** 10.1101/2025.04.11.648468

**Authors:** Da Zhang, Jonathan K. Kleen

**Affiliations:** Department of Neurology, University of California San Francisco, San Francisco, CA 94143; Weill Institute for Neurosciences, University of California, San Francisco, CA, 94158

## Abstract

Two frequently studied bursts of neural activity in the hippocampus are normal physiological ripples and abnormal interictal epileptiform discharges (IEDs). While they are different waveforms, IEDs are notoriously picked up as false positives when using typical automated ripples detectors which are prone to sharp edge artifacts. This has created challenges for studying ripples and IEDs independently. We leveraged recent advances in computer vision on time-frequency feature representations to enable more comprehensive and objective dissociation of these phenomena. We retrospectively evaluated human intracranial recordings from 46 hippocampal depth electrode sites among 17 patients with focal epilepsy, the majority of whom had a seizure-onset zone/network involving the hippocampus. We implemented a common human ripple detection algorithm and broadband spectrograms of all detected “ripple candidates” were projected into low-dimensional space. We segmented them using k-means to infer pseudo-labels for probable ripples and probable IEDs. Independently, human expert IED labels were manually annotated for comparison. State-of-the-art vision transformer models were implemented on individual spectrograms to approach ripple vs. IED dissociation as an image classification problem. We detected 31,847 ripple/IED candidates, and a median 3.9% per patient (range: 0-47.2%) were IEDs based on expert label overlap. Low-dimensional projection of spectrograms separated canonical IEDs vs. ripples better than raw or ripple-filtered waveforms. Canonical ripple and IED candidates emerged at opposite poles with a continuous landscape of intermediates in between. A binary vision transformer model trained on expert-labeled IED vs. non-IED candidate spectrograms with 5-fold cross-validation showed a mean area under the curve (AUC) of 0.970 and mean precision-recall curve of 0.694, both significantly above chance. To evaluate generalizability, we implemented a leave-one-patient-out cross-validation approach, in which training on pseudo-labels and testing on expert-labeled data demonstrated near-expert performance (mean AUC 0.966 across patients, range 0.892-0.997). Transformer-derived attention maps revealed that models were tuned to triangle-like edge artifact spatial features in the spectrograms. Model-derived probabilities (i.e. of being an IED) for all candidates demonstrated continuous transitions between ripples vs. IEDs, as opposed to binary clustering. The delineation between ripples and IEDs appears best represented as a gradient (i.e. not binary) due to physiological ripple features overlapping with sharpened and/or high frequency pathophysiological IED features. Vision transformers nevertheless perform virtually at human expert levels in dissociating these phenomena by leveraging time-frequency spatial features enabled by neural data spectrograms. Such tools applied to spectrotemporal representations may augment comprehensive investigations in cognitive neurophysiology and epileptiform signal biomarker optimization for closed-loop applications.

## Introduction

The hippocampus is a key brain region for memory function, including remembering what we did minutes or even decades ago.^1^ Extensive preceding rodent literature has described the morphological, spectral, and neurophysiological features of transient events called “ripples” (**Fig. 1A-C**) that are dynamic physiological signatures of hippocampal memory processing.^2^ Hippocampal ripples are brief (∼100ms) bursts of hippocampal activity occurring in the high gamma range frequency range (110-180 Hz range, or 70-150 Hz in humans)^3^ generated from intricate interactions between trilaminar CA3 and CA1 subfields in hippocampal circuits.^2^ They are often termed sharp-wave ripples since, as demonstrated in animal models and variably in humans,^2,3^ they can be superimposed upon delta waveform transients (of note, this is distinct from pathophysiological scalp electroencephalogram “sharp waves” described in epilepsy^4^). Hippocampal ripples are demonstrated across several species and they play a major role in encoding, consolidation, and even real-time memory retrieval described more recently in humans.^1,2,5,6^

**Figure 1.**
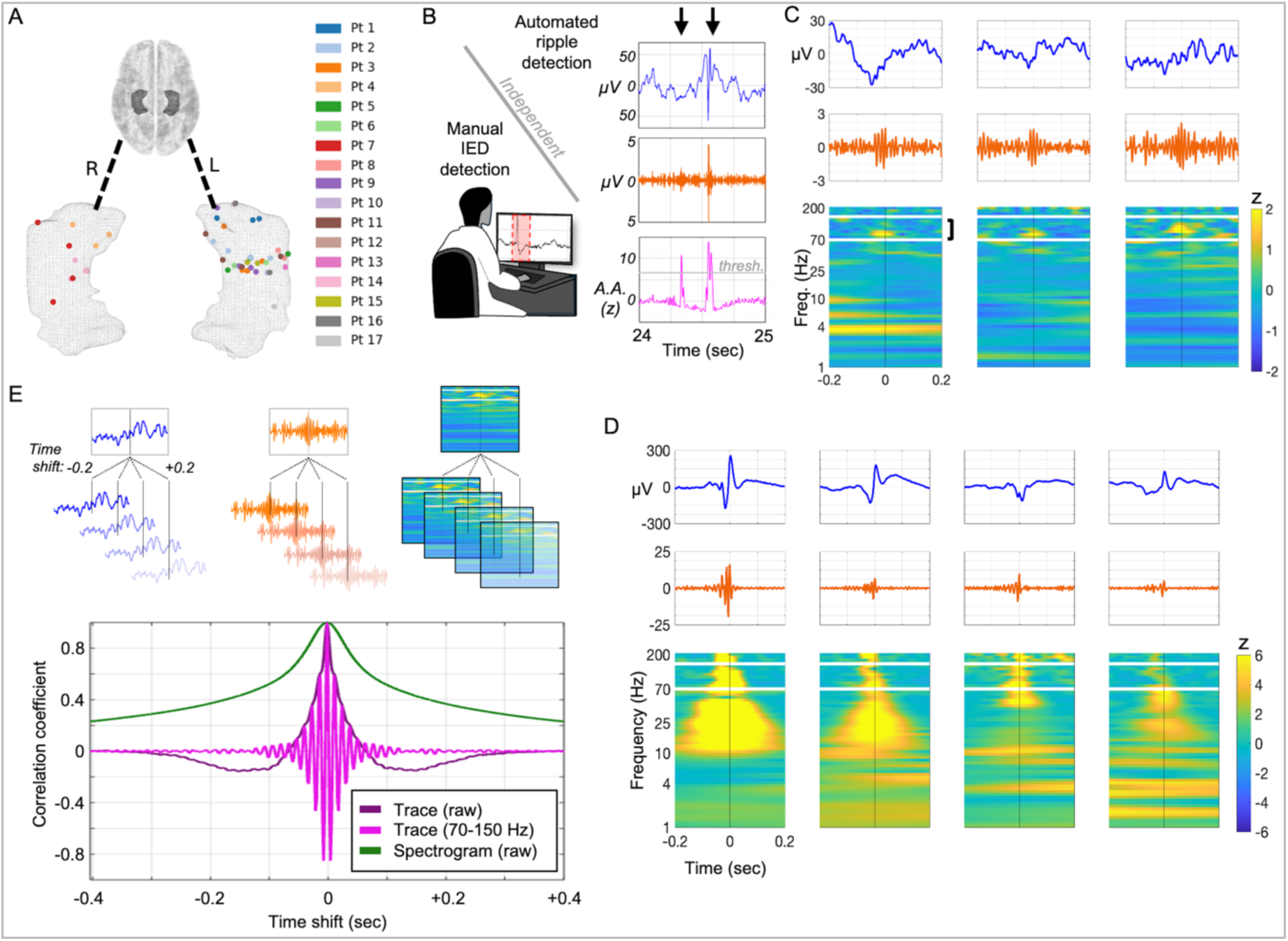
Hippocampal ripple and IED time-frequency profiles. **A**. Hippocampal depth electrode coverage across all patients, illustrated as approximate positioning of electrode contacts for each patient (colors) warped to average reconstructions of right and left hippocampi (inferior view, shown within 3D glass brain at top, reconstructions using MNI152) **B.** Conceptual schematic of manual expert label annotation (bottom-left panel) which was done independently of automated ripple/IED candidate detection. Right-sided panels demonstrate raw voltage trace (top), ripple-filtered (70-150 Hz; middle), and z-scored analytic amplitude (bottom; gray line: example detection threshold). Left and right black arrows show detected ripple and IED respectively. **C.** Examples of detected candidates (panel columns) consistent with normal physiological ripples (raw trace, ripple-filtered trace, and spectrograms in top, middle, and bottom rows). time-aligned to phase trough nearest to maximal analytic amplitude of ripple frequency band (70-150 Hz; also outlined by black bracket and white lines in spectrograms. Note subjective difficulty in visualizing ripple oscillation (middle) in raw data (top). **D.** Examples of detected candidates (panel columns) expert-labeled as IEDs with more subtle epileptiform features moving towards the right. Note general similarity of (i.e. subjective difficulty in distinguishing) IED traces filtered in the ripple band (middle panels) versus normal physiological ripples (middle panels in C). **E.** Time-shifted autocorrelation of the raw trace filtered trace, and spectrogram (1–200 Hz) data centered window (-0.1 through +0.1 seconds) computed over signal lags ranging from -0.4 s to 0.4 s. Data beyond center window utilized but not shown in schematic at top. Inversions of raw and ripple-filtered traces likely due to anti-phase correlations of low and ripple-band frequencies prominent in these respective signals.

Human hippocampal ripples are recorded using intracranial electroencephalogram recordings (ICEEG) in patients with drug resistant seizures who are undergoing monitoring for epilepsy surgery to localize the seizure-onset zone (SOZ).^1,3,6^ During ICEEG recordings, brain regions overlapping with the SOZ, which commonly include the hippocampus,^7^ often also generate brief spontaneous bursts of pathological electrical activity called interictal epileptiform discharges (IEDs). IEDs can dynamically commandeer and disorganize ongoing neural representations of information,^8^ conferring transient cognitive improvements with cumulative influences.^9–11^. The combination of static baseline impairments (e.g. due to hippocampal sclerosis) and dynamic influences (IEDs and seizures) leads to the cognitive impairments pervasively affecting daily life and even scholastic performance.^12,13^

IEDs and physiological ripples are frequently observed in the same hippocampal electrode sites^3,9,14^ because of abnormal excitatory connectivity, reduction of inhibitory tone, and other circuit changes.^2,15^ Their anatomical overlap is further complicated by similarity in duration (generally <200ms), timing (sporadic), and importantly, signal features (**Fig. 1C,D**). Specifically, physiological ripples have variation in their spectral signatures including peak frequency, spectral bandwidth, and lower frequency admixtures manifesting the “sharp wave” component and even the current behavioral state.^3,16^ IEDs can mimic ripple features due to their “sharp” or “spiky” waveform morphology^4,9^ producing sharp edge-related high frequency artifact,^17,18^ and superimposed high-frequency oscillations (HFOs, 50-500 Hz).^19–21^

A variety of ripple and IED detectors are used frequently in the field, as manual review of resultant high volumes of detected events across long recordings is not feasible.^6,9,10,18^ Yet IEDs pervasively produce false positives by triggering standard ripple detectors^18^ which almost exclusively rely on frequency-based parameters.^3^ The challenge of distinguishing physiological ripples from IEDs in human cognitive neurophysiology rests largely with this blurred overlap in frequencies.^3,19^ A landmark paper by Engel *et al* in 2009^19^ stated that “frequency alone is not sufficient for separating normal from pathologic oscillations”. Nevertheless, detected human hippocampal ripples are often assumed as distinct despite potential IED contamination among variable proportion of the detections. Manual screening is considered the gold standard for verification of both ripples^3^ and IEDs,^9,18^ however subjectivity and inter-rater reliability undermine this key quality check, especially for ambiguous or borderline epileptiform features.^3,22,23^

Importantly, only a small fraction of detected ripple candidates may be true hippocampal ripples. For example, one recent study using a refined combination of objective and subjective criteria demonstrated that only 11% of ripple candidates gathered after automated ripple detection were felt to be consistent with true physiological ripples.^16^ This again underscores the unclear nature and the physiological roles of most other detected ripple-like events in humans. The spectrotemporal heterogeneity of ripples, IEDs, and other ripple-like and IED-like candidates has substantial relevance for understanding hippocampal function and dysfunction,^3,9,10,24^ as well as improving biomarker optimization (e.g. IED and/or seizure detection) for closed-loop neurostimulation treatments.^25^ Traditional binary (ripple vs. IED) categorization is agnostic to these potentially rich details.

Here we addressed these challenges by 1) avoiding assumptions that ripples and IEDs are always distinct, and 2) leveraging their spectrotemporal (i.e. spatial, two-dimensional) features using recent advances in computer vision and artificial intelligence (AI). Specifically, we implemented low-dimensional embeddings of broadband spectrogram data, and applied vision transformers (ViT) for spectrogram classification as an image classification problem. The latter also provided information on crucial data features via the ViT attention-based mechanism (as opposed to traditional convolutional networks in which key features are hidden in deep layers). We hypothesized that hippocampal ripples and IEDs exist along a continuum of spectrotemporal gradients that provides a novel framework not only dissociating them but also evaluating their heterogeneity through quantitative means.

## Methods

### Participants and data acquisition

We studied the intracranial recordings of seventeen patient participants with hippocampal electrode coverage who were undergoing intracranial monitoring for drug-resistant epilepsy at our institution (Table 1). Written informed consent was obtained for all participants, and this work was approved by the UCSF Institutional Review Board.

**Table 1.** Patient characteristics. Abbreviations: L: left, IFG: inferior frontal gyrus, ITG: inferior temporal gyrus, MFG: middle frontal gyrus, MTG: middle temporal gyrus, STG: superior temporal gyrus, PVNH: periventricular nodular heterotopia. *Based on interictal findings due to no recorded seizures.

| ID | Age | Implant side | Gender | Seizure-onset zone/network |
| --- | --- | --- | --- | --- |
| Pt 1 | 37 | Left | Female | Anterior/mesial/basal temporal |
| Pt 2 | 47 | Left | Female | Posterior lateral temporal region |
| Pt 3 | 36 | Left | Male | Mesial temporal |
| Pt 4 | 38 | Bilateral | Male | Mesial temporal* |
| Pt 5 | 51 | Left | Female | Mesial temporal |
| Pt 6 | 38 | Left | Male | Mesial temporal |
| Pt 7 | 36 | Bilateral | Female | Lateral temporal and frontal (multifocal) |
| Pt 8 | 37 | Left | Male | Anterior/mesial temporal |
| Pt 9 | 19 | Left | Male | Posterior MTG, STG |
| Pt 10 | 21 | Left | Female | Inferior lateral temporal |
| Pt 11 | 22 | Left | Female | Posterior lateral temporal and frontal (multifocal) |
| Pt 12 | 43 | Left | Male | Superior frontal encephalomalacia |
| Pt 13 | 21 | Left | Male | Posterior STG |
| Pt 14 | 20 | Right | Male | Mesial temporal |
| Pt 15 | 33 | Left | Male | Lateral occipito-temporal junction |
| Pt 16 | 24 | Left | Female | Periventricular nodular heterotopia |
| Pt 17 | 49 | Left | Female | Mesial temporal |

Hippocampal signals were recorded from linear depth electrode probes implanted in either or both hemispheres. We verified hippocampal recording sites for each patient by co-registering their preoperative T1 MRI with electrode coordinates from their postoperative CT scan using the SPM12 software package (SPM12, https://www.fil.ion.ucl.ac.uk/spm/software/spm12). Electrodes were then classified to the hippocampal regions based on visual inspection of the co-registered images, and we plotted hippocampal electrode coverage across all patients using patient-specific nonlinear surface registration aligned in Freesurfer and warped to an average brain volume template (CVS atlas in MNI152 space; **Fig. 1A**).^26–29^ Following verification of hippocampal positioning and inspection of raw recordings, channels with good signal quality were selected for analysis, with a maximum of four electrodes for each patient.

Neural data voltage signals were recorded at a sampling rate of 3052 Hz on a multichannel amplifier optically connected to a digital signal processor (Tucker-Davis Technologies Inc., Alachua, FL) in awake patient participants alternating between rest and performance of a verbal memory task implicating the hippocampus and medial temporal lobe (adapted auditory naming).^9,30^ Signals were visually examined and any channels containing low signal, electrical noise, or other substantial artifacts were excluded from future analysis, since these factors can cause artifactual contamination of both ripple and IED analysis.^18^ An anti-aliasing filter was applied and the neural signals were then downsampled to 512 Hz and notch-filtered (60 Hz and harmonics) to remove line noise.

### Manual IED annotation

The voltage data traces from the hippocampal ICEEG recordings were manually annotated by a board-certified epileptologist (J.K.K.) by plotting on a custom graphical user interface developed in MATLAB (Natick, MA). All recorded channels were displayed simultaneously to provide anatomical context (e.g. overall field, propagation, etc.) during interpretation. The duration and specific channels involved in an IED were annotated by first using a linelength-based automated detector^31^ followed by manual screening and correction for false positives and negatives.^18^ Importantly, all manual annotation (expert labeling) procedures above were prior to, and independent of, the ripple candidate detection procedures (**Fig. 1B**).

### Candidate detection and signal processing

We detected ripple/IED candidates on these hippocampal electrode recordings using established threshold-based methods^3,6^ that were adapted by removing IED screening steps. Specifically, along the approaches of Norman *et al*.^6^ for each recording session, we converted the raw ICEEG signal to continuous analytic amplitude (AA) in the 70-150 Hz bandpass range. These 70-150 Hz AA signals were z-scored for each frequency and for each recording block. We then clipped the data at 4 standard deviations (SD) to help mitigate data skewing, squared the data, then smoothed the resulting signal using a 1-40 Hz bandpass Butterworth filter. The mean and SD of this resulting signal were computed, and timepoints from the original squared yet unclipped signal that were 4 SD above the mean of this signal were considered preliminary detection candidate events. Any candidates less than 30ms from each other were merged as one candidate. The durations of ripple candidates were computed from the first timepoints above the 4 SD threshold through the timepoint dropping below 2 SD, and events with durations <20MS or >200ms were excluded. Candidates were re-aligned to the timepoint of the ripple oscillation waveform trough nearest to the peak ripple amplitude.^1,6^

IEDs were naturally included among the detections (**Fig. 1B**, right panels) due to the high-frequency signals generated by sharp edges (spectral leakage)^18^ and pathological high-frequency activity inherent to epileptiform waveforms. Numerous publications describe various methods of identifying and excluding epileptiform detections from ripple datasets nicely summarized by Liu et al,^3^ however there is no current universally accepted technical standard nor formalized training for such screening procedures – instead, we waived IED exclusion steps since the issue of overlap in ripple and IED detection is the crux of our study (**Fig. 2**). This allowed us to then account for their hypothesized heterogeneity along a normal-abnormal gradient space through low-dimensional data reductions,^3,32^ and moreover, to test whether they are nevertheless dissociable by leveraging novel spatial feature analysis of their spectrotemporal representations with ViT-based classification.

**Figure 2.**
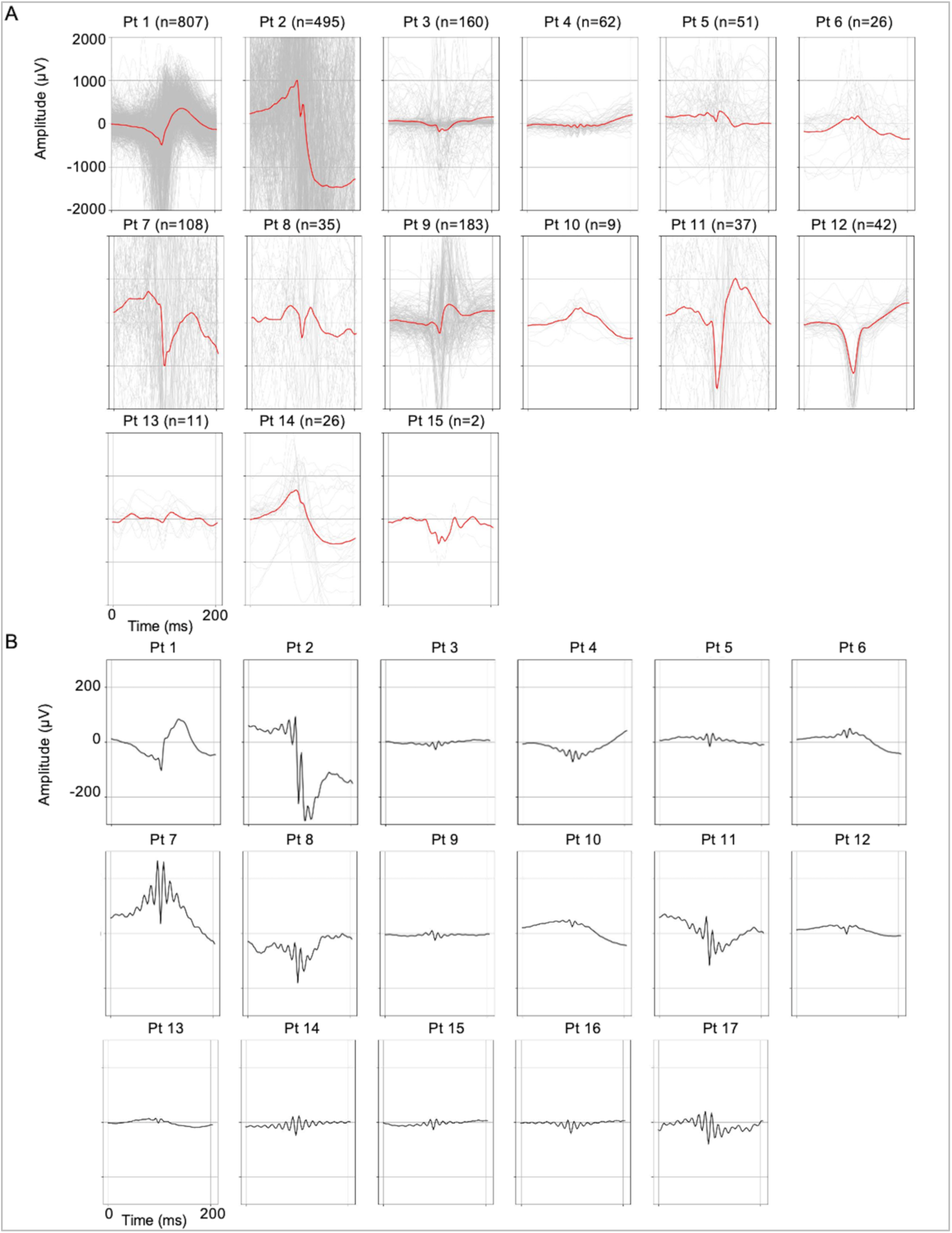
Detected IED and non-IED candidates across all patients. **A**. Raw traces (gray) and their average (red) for detected candidates that were independently expert-labeled as IEDs, for each patient. Wide heterogeneity in epileptiform (sharp and spiky) features can be observed both between and within patients. Patients 16 and 17 had no expert-labeled IEDs. **B**. Average traces for detected candidates that were not expert-labeled as IEDs (non-IEDs) for each patient, demonstrating superimposed delta wave components for some, yet also sharpened aspects likely related to epileptiform contamination and reflecting borderline/gray zone candidates ultimately not marked manually.

For all candidates, we stored three representations (substrates) using ±0.1 sec from center timepoints at 512 Hz sampling rate for a total of 103 timepoints. These were raw waveforms (1-D vector), ripple-filtered waveforms (70-150 Hz, using a Butterworth band-pass filter; 1-D vector), and spectrograms (2-D matrix). We extracted these latter peri-event broadband spectrograms by converted the raw continuous ICEEG signal from each block to continuous Hilbert transform AA for 61 logarithmically spaced frequencies in the 1-200 Hz range. These signals were z-scored for each frequency, and for each recording session. These three candidate data substrates are represented in the respective top, middle, and bottom panels of **Fig. 1C,D**).

### Temporal autocorrelations

Ripples and IEDs vary widely in absolute voltage at each time point (examples in top panels in **Fig. 1C,D**). Alignment to these variable waveform characteristics (e.g. deciding between alignment to raw waveform peak vs. trough vs. maximal slope) imparts substantial timepoint-by-timepoint variation in downstream calculations. This is relevant for filtered signals as well, since ripple band oscillations rapidly change voltage sign within a few timepoints (examples in middle panels of **Fig. 1C,D**). These aspects could therefore influence or potentially undermine clustering and classification when using raw waveforms, and thus we compared between raw waveform, ripple-filtered waveform, and spectrogram data substrates using signal autocorrelations to assess the impact of small time shifts compared to raw voltage (**Fig. 1E**). Specifically, we use the center 200ms of each candidate autocorrelated with the same signal lagged from -0.4s - 0.4s, implementing the autocorrelation formula *Corr*(*τ*) = (*x*_*t*_, *x*_*t*"*τ*_), where *Corr*(*τ*) is the Pearson-based autocorrelation at lag *τ*, and *x*_*t*_ is the value of the signal at time *t*.

### Dimensionality reduction

We applied uniform manifold approximation and projection (UMAP) as a nonlinear dimensionality technique to reduce and visualize the vectorized candidate spectrograms into low-dimensional.^32,33^ We fixed UMAP parameters in the ranges of prior work,^16,32^ rather than iterating over a wide parameter space, since the central objectives of this analysis were 1) to assess embedding distributions using standard (off-the-shelf) parameters, and moreover 2) to compare between data substrate conditions (raw trace, ripple-filtered trace, and spectrogram; **Fig. 3A-C**). Specifically, we used a Euclidean metric and set the number of components as *n_components=2*, the number of neighbors as *n_neighbors=15*, and the minimum distance as *min_dist=0.1*. Following the original UMAP run using a random seed, we fixed this parameter (*random_state=42*) for replicability. To segment putative subpopulations of the detection candidates into an inherent groups in an objective (unsupervised) manner, we then employed K-means as an clustering approach using the first two UMAP dimensions.^34^ To identify the optimum number of clusters we conducted a hyperparameter search ranging from K = 3 to 20 and evaluated silhouette scores for the resultant segmented clusters (**Fig. 3E)**.

**Figure 3.**
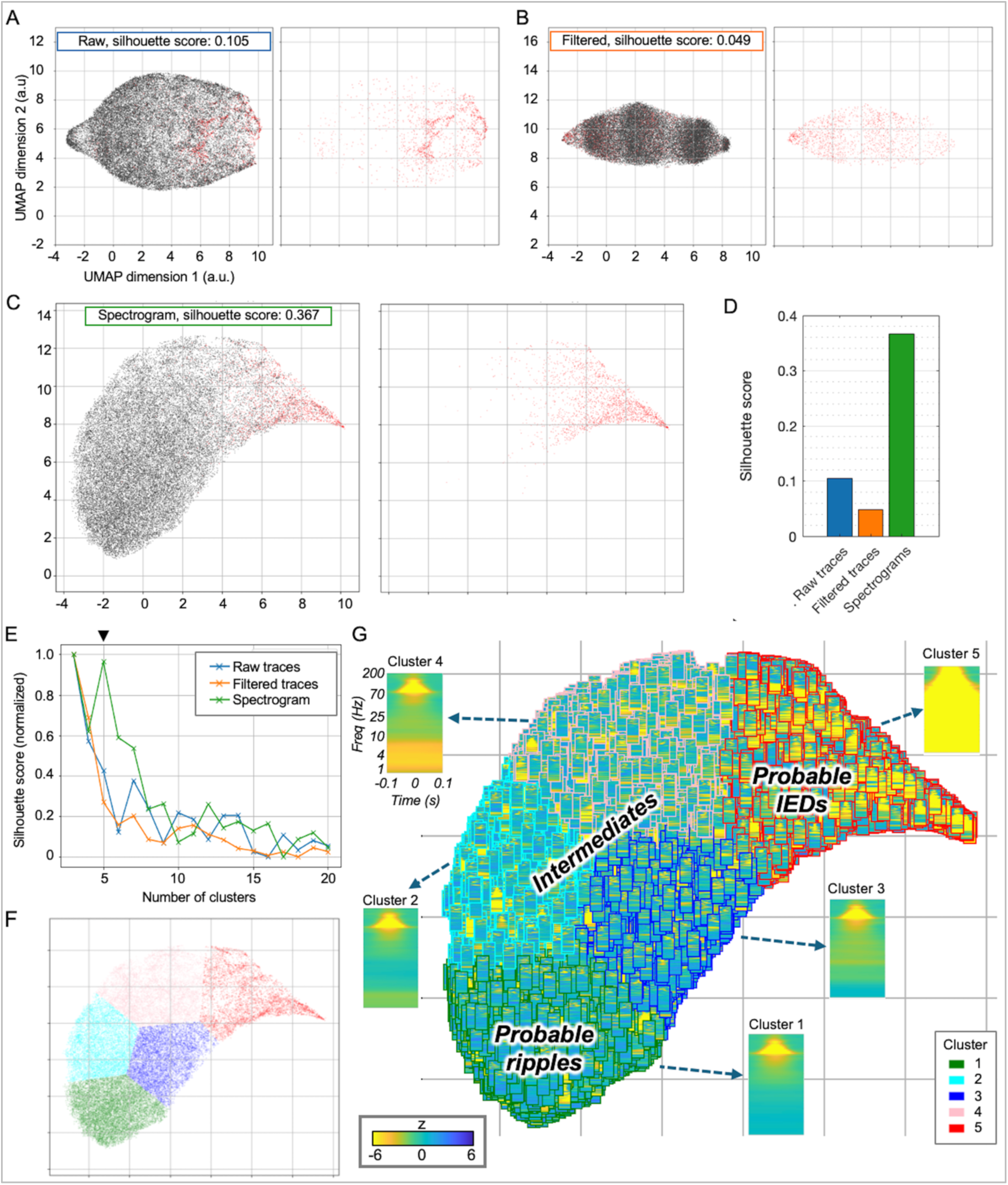
Low-dimensional projection, clustering, and segmentation of detected candidates. **A.** All detected candidates across all patients (left panel) plotted in low-dimensional embedding (first two UMAP dimensions) using raw voltage traces (examples in top panels of Fig. 1C,D). Expert-labeled IEDs are shown as red dots (isolated in right panel) whereas all other candidates are black dots. Silhouette score at top. **B.** Same as A using ripple-filtered traces (70-150 Hz; examples in middle panels of Fig. 1C,D. **C.** Same as A and B using spectrograms (examples in bottom panels of Fig. 1C,D). **D.** Supervised label silhouette scores for A-C using two groups as binary labels (expert-labeled IEDs vs. all other candidates). **E.** Unsupervised cluster label silhouette scores derived from k-means iterated over 3 through 20 clusters (normalized to scores at n=3 clusters) for raw trace, ripple-filtered trace, and spectrogram UMAP distributions (left panels of A-C). A secondary peak is illustrated at n=5 clusters for spectrogram data (black arrow), which was thus used for subsequent unsupervised segmentation of all candidates into candidate groups for further analysis. Note the lack of a comparable secondary peak in raw and ripple-filtered data. **F.** Visualization of the five unsupervised spectrogram-based cluster labels (colors) derived from E on the corresponding spectrogram UMAP embedding from C. **G.** Segmented data points from F visualized with individual spectrograms in corresponding embedding positions, as well as average spectrograms for the five clusters. Candidates in Cluster 5 were henceforth considered probable IEDs in light of strongest overlap with expert-labeled IEDs in C, whereas candidates in Cluster 1 were considered probable ripples in light of polar opposite position. Clusters 2-4 were considered “intermediates”, in light of gradient-like UMAP distributions of detected candidates (i.e. lack of clear separable, isolated clusters) regardless of data type.

To ensure the different regions of the low-dimensional distribution were not driven by a limited number of patients, we divided the UMAP space into 20 sections in both the X and Y dimensions (20×20; **Fig. 4**) and calculated the Shannon entropy values^35^ based on the equation:

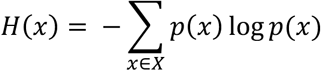

**Figure 4.**
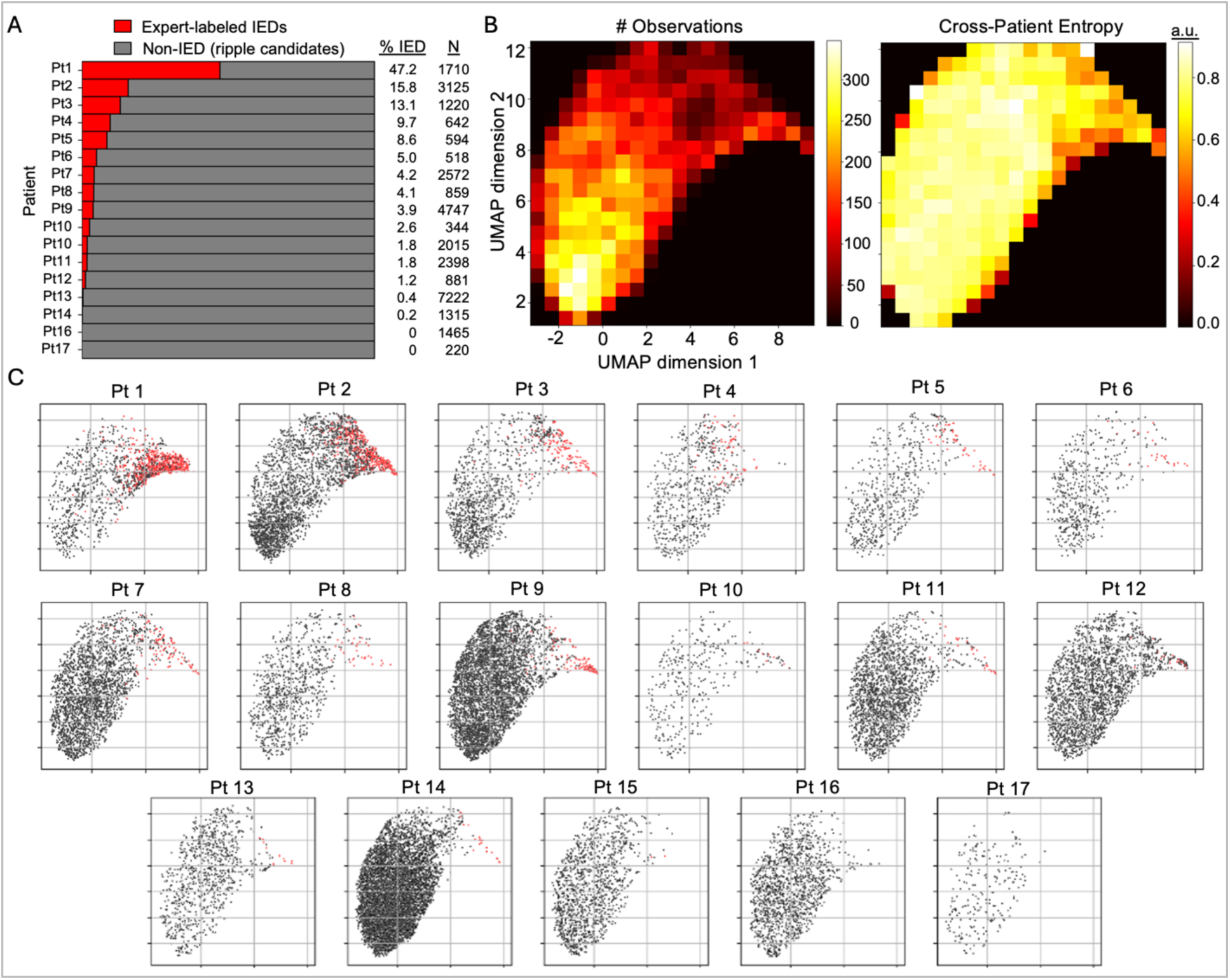
Individual-level distributions of detected candidates. **A.** Proportion of expert-labeled IEDs (red) and non-IEDs (gray) for each patient, sorted by the IED proportion. **B.** Density of number of observations in UMAP space (see Fig. 3) across patients (left panel), and cross-patient entropy demonstrating relatively similar contribution of different patients to nearly all regions of this same distribution (right panel). **C.** Patient-level distributions of expert-labeled IED and non-IED candidates (similar to plot for all patients in Fig. 3C), demonstrating relatively similar positioning of expert-labeled IEDs in upper-right region UMAP space.

Here, *x* represents individual samples for each patient and *H*(*x*) quantifies the entropy value, representing the uncertainty or information content within the distribution. The term *p*(*x*) represents the probability mass function for *x*, indicating the relative frequency of samples within each division. This provided a mathematical framework to assess the degree of randomness or heterogeneity of the information, where higher entropy values correspond to greater variability in the patients contributing to a given region of the UMAP embedding.

### Vision transformer models

A power spectrogram provides a comprehensive representation of information contained in a time series signal (each frequency at each timepoint); minimal information is lost when converting back-and-forth between a raw signal and its spectrogram. Importantly, humans can evaluate this two-dimensional matrix layout of data by eye, as an image (i.e. like viewing a picture). Consider a photo of a dog – we recognize it because the ears are located in specific places relative to the snout, mouth, legs, etc. (e.g. the spatial features, or relative layout, of the dog’s key parts in the photo; **Fig. 5A**). Similar to how we interpret the spatial relational features of photos by pixels arranged in columns and rows, spectrogram features can be analogously interpreted in time and frequency axes (**Fig. 5A**).

**Figure 5.**
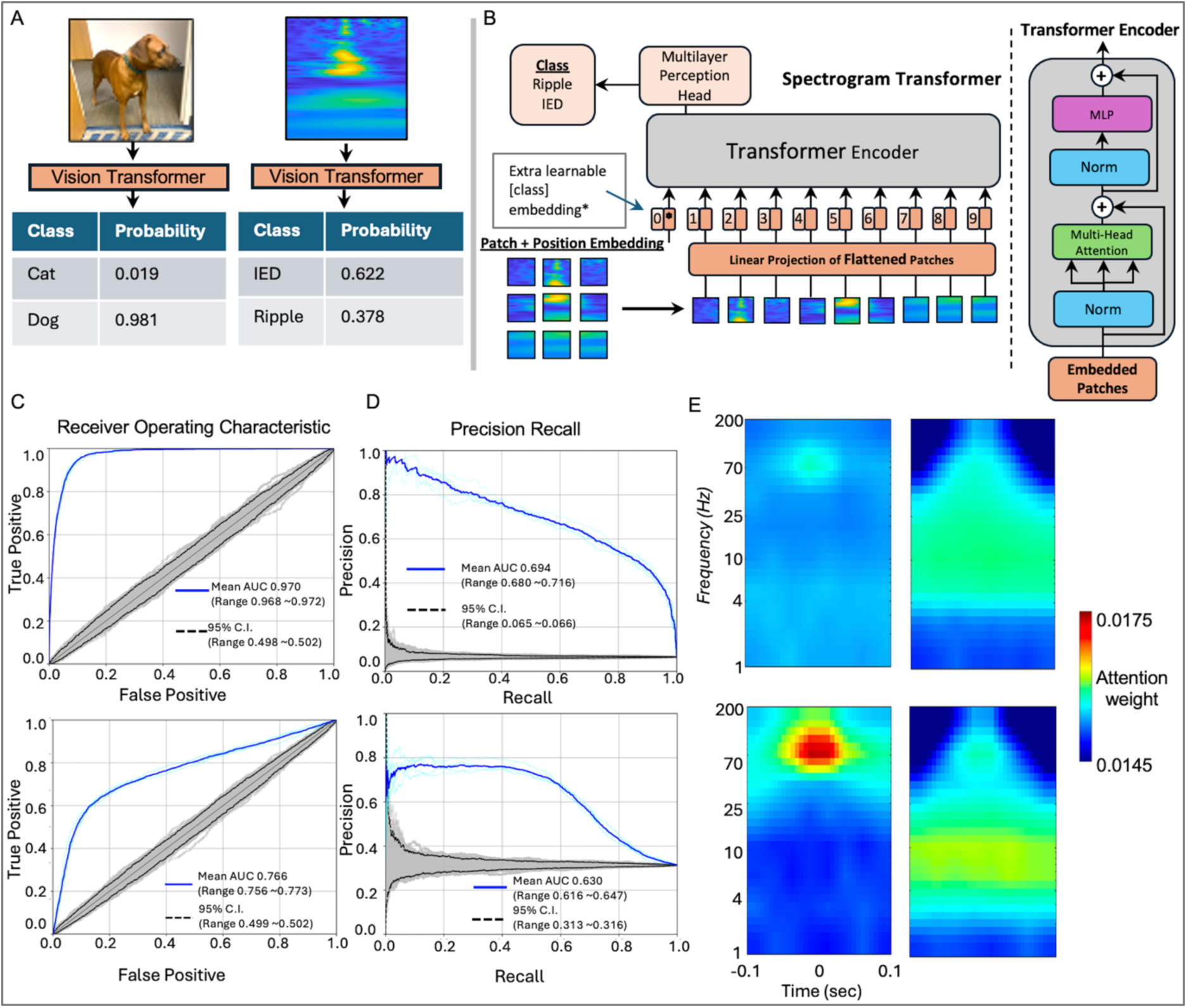
Spectrogram vision transformer for ripple/IED detection. **A.** Concept schematic of ViT image classification mechanism, applied to a photo to discern the probability of being a dog (left panels; probabilities derived from a ViT model^42^ pre-trained on ImageNet-21k and ImageNet 2012), and in our method applied to a spectrogram to discern the probability of being an IED (right panels; trained on our data as a binary image classification problem). **B.** Architecture of SG-Transformer model demonstrating patch and position embedding, thereby preserving the spatial structure of the spectrogram data (relative time and frequency relational information). **C.** AUC-ROC scores for 5-fold cross-validated SG-Transformer binary classification model (i.e. probability of being an IED; blue trace and envelope: mean ±SEM of 5 folds), using expert-derived labels (IEDs vs. non-IEDs; top panel) and pseudo-labels from probable ripples and probable IEDs (Clusters 1 vs. 5, refer to Fig. 3F,G; bottom panel). Both far exceeded the 95% confidence interval (black outline) from 200 shuffled iterations (gray traces). **D.** PR-AUC scores from same models as in C, again far exceeding 95% confidence intervals. **E.** Attention maps for non-IEDs and probable ripples (respective top and bottom panels on left) and expert-labeled IEDs/probable IEDs (and probable ripples (respective top and bottom panels on left) corresponding to the models in C and D.

To identify the signal features most relevant for dissociating ripples and IEDs, we adapted a ViT model to leverage the spatial relationships of time-frequency features in spectrograms (i.e. interpreting spectrograms as images). This model architecture, which we call SG-Transformer (Spectrogram-Transformer), relied upon the state-of-the-art “attention” mechanism in image classification^36^ applied to the two-dimensional data of a spectrogram, as alluded to in recent scalp EEG work^37–40^ though to our knowledge not yet applied in ICEEG.

The SG-Transformer was trained as a binary classifier to distinguish between the ripples and IEDs using their 200ms peri-event spectrograms (see above) as data substrates. Unlike convolutional neural networks (CNNs) or time series-based networks (e.g. RNN, LSTMs^41^), which primarily capture the local perceptive field, the attention-based ViT model allowed patches within an image to interact with each other through its self-attention mechanism, enabling the capture of global (spatial) information present in the spectrogram. In other words, each spectrogram is treated as an image and then as a sequence of patches, and each patch is fed into the linear projection first and then passed through the encoder model, similar to image classification applications^36,42^ (**Fig. 5A,B**).

The SG-Transformer model was adapted from the original ViT architecture repository by Dosovitskiy *et al*.^42^ We used a two-layer multi-head self-attention module and set the number of heads to 2 for each layer. During training, we use an AdamW^43^ optimizer with an initial learning rate of 10^−5^ and a weight decay of 10^−5^ with the batch size setting to 256. Models were cross-validated using five folds (80% training data, 20% testing data) and later by a leave-one-patient-out (LOPO) approach. All models (folds) were trained for 100 epochs each, and the epoch with the optimal performance (smallest loss) on the test set was used. Models were trained on two NVIDIA RTX A5000 GPUs. We used a receiver operating characteristic curve (AUC-ROC) for prediction performance metrics. Additionally, since the dataset is imbalanced between classes with more non-IED candidates than IED candidates, we also implemented precision-recall area under the curve (PRAUC) analyses. The chance levels for PRAUC, (i.e. the value obtained after randomly shuffling the dataset), was calculated as the number of positives samples divided by the total number of samples.

## RESULTS

### Patient characteristics

We studied the hippocampal depth electrode recordings of seventeen patient participants (14 left-sided, 1 right-side, 2 bilateral) who were undergoing intracranial monitoring for drug-resistant epilepsy at our institution (**Table 1**). Ages ranged from 19-51 years, and 47% identified as female and 53% identified as male. Nine of the patients (52.9%) had confirmed mesial temporal and/or hippocampal involvement in the seizure-onset zone, and 15 of 17 patients (88.2%) had hippocampal IEDs based on manual expert review (**Fig. 4A**).

### Candidate detections

We implemented an automated detector targeting physiological ripples based on Norman *et al*.^6^ (see Methods), resulting in 31,847 detections across the 17 patients, which we refer to henceforth as ripple “candidates.” The timestamps of these candidates were cross-checked against the manual IED detections, and candidates with timing that overlapped with manual detection were marked as IED detections. Initial screening of the candidates revealed numerous obvious ripples (**Fig. 1C**), but also many IEDs, along with intermediate events that had variable degrees of both ripple-like and IED-like features and abnormal features as anticipated (see Methods)^18^, examples of which are shown in **Fig. 1C,D**.

Expert-labeled IEDs among the candidates ranged from 0.2% to 47.2% of the detections per patient (**Fig. 2A**), for a total of 2,054 IEDs across all patients. Inspection revealed variable morphologies similar to prior descriptions^31^ these were generally distinct (i.e., largely stereotyped) for each patient (**Fig. 2A**) as anticipated. We also examined all other detected waveforms (putative ripples) filtered in the original detection band (70-150*** Hz).

We next performed a comparison of time-shifted autocorrelations for raw waveforms, ripple-waveforms, and spectrogram signal representations (**Fig. 1C,D**). Raw waveforms demonstrated a relatively fast drop-off in autocorrelation values, within roughly ± 50ms of the centered signal, and inverse autocorrelation values within roughly ± 50-250 ms likely related to the low-frequency components. Ripple-filtered waveforms showed extremely fast drop-offs and oscillating signal inversions at large correlation extremes that tapered off by ± 100ms reflecting of their oscillatory features within expected ripple candidate durations (<200ms total). Spectrograms demonstrated comparatively greater temporal stability (reduced sensitivity to time shifts) with a gradual drop-off over hundreds of milliseconds. The resultant two-dimensional (61 x 103) spectrogram matrices were used as the data substrate for all further analyses.

### Low-dimensional analysis

We applied UMAP to the 31,847 candidates as raw waveform and ripple-filtered waveforms and projected them into an embedding space (**Fig. 3A,B**). Each demonstrated a relatively homogenous main cluster of data with relatively wide overlap of expert-labeled IEDs vs. non-IEDs when viewed visually, confirmed quantitatively by silhouette scores of 0.105 (raw; **Fig. 3A**) and 0.049 (ripple-filtered; **Fig. 3B**). We then applied UMAP to vectorized candidate spectrograms, again demonstrating a relatively homogeneous-appearing large oval cluster of data, yet with a tapered end (top-right regions of panels in **Fig. 3C**) which was more concentrated with expert-labeled IEDs and a substantially higher silhouette score of 0.367 (**Fig. 3D**).

Since the data distributions in UMAP space demonstrated the hypothesized wide gradients (as opposed to clear separable clusters) between expert-labeled IED and non-IED candidates which may have physiological-pathophysiological relevance, we then evaluated unsupervised clustering of the data to derive pseudo-labels through data segmentation using k-means. When iterating over 3 through 20 possible clusters derived from k-means, we found steep drop-offs in silhouette scores for raw and ripple-filtered waveforms. Spectrograms on the other hand demonstrated a secondary peak in the quality of clustering at k=5 clusters (**Fig. 3E**). This suggested roughly optimal segmentation of the data in UMAP space and thus we henceforth used this cluster ID information as pseudo-labels for the data in the remainder of the study (**Fig. 3F**). Average spectrograms for all candidates in each cluster are illustrated in **Fig. 3G**. The UMAP cluster overlapping most with expert IED labels was cluster 5 (**Fig. 3 C,F,G**); this and the cluster at the opposite pole of the projection (cluster 1) were thus considered as best representative of canonical IEDs and canonical ripples, respectively, underscored by their spectrogram appearances (e.g. compare spectrograms **Fig. 3G** vs. examples in **Fig. 1B,C**).

### Participant-level distributions

In our dataset 15 of 17 of patients (88.2%) had detected candidates consistent with expert-labeled IEDs (Pt 16 and Pt 17 had no expert-labeled IEDs detected in their data). Across these patients, the percentage of these IED candidates was skewed widely from 0.02 to 47.2% with a median of 4.1% (**Fig. 4A**). We plotted the binned counts of the number of candidates in each window, demonstrating a higher density of candidates in the ripple-like end of the UMAP distribution (**Fig. 4B**, lower right region of left panel). To ensure generalizability (i.e. that no single participant predominantly drove data distributions), we plotted the cross-patient entropy for each section (**Fig. 4B**, right panel). This was relatively consistent across the UMAP embedding suggesting of boundary effects. Underscoring this idea further, projecting individual patient data in this same space (**Fig. 4C**) demonstrated relatively similar candidate embeddings widely across the breadth of the distribution. Moreover, expert-labeled IEDs tended to fall in the same upper right region of the distribution (region of cluster 5), suggesting consistency when using spectrograms in this scenario despite the variety of IED morphologies across patients (**Fig. 2A**).

### Dissociating ripples and IEDs

The results in **Fig. 4** demonstrate that distributions of putative ripples and putative IEDs localized consistently across patients to relatively opposite ends (poles) of the overall distribution, and we used this framework to create unsupervised labels using the k-means clusters corresponding to putative ripples (Cluster 1) and putative IEDs (Cluster 5). We implemented our SG-Transformer essentially as a binary image classification model on the two-dimensional spectrograms (**Fig. 5A,B**) by evaluating classification performance with a 5-fold cross-validation method.

We initially trained the model on candidate spectrograms using expert labels (IED vs. not IED), which demonstrated a mean AUC-ROC score across the five folds of 0.970 (range: 0.968 to 0.972) and a PRAUC mean across folds of 0.694±0.012 (range 0.680 to 0.716). These values were well beyond the 95% confidence interval of 200 iterations in which the IED and ripple labels were randomly shuffled (**Fig. 5C,D**; 95% C.I.: 0.498-0.502).

We next trained the SG-Transformer classifier model on pseudo-labels (probable ripples vs. probable IEDs, i.e. Clusters 1 vs. 5, respectively). The AUC-ROC score across the five folds ranged from 0.756 to 0.773 with a mean of 0.766±0.006 (**Fig. 5C**, lower panel), beyond chance levels (95% C.I. 0.499-0.502). The mean PRAUC across folds was 0.630±0.012 (range: 0.616 to 0.647), again above chance (95% C.I.: 0.313 to 0.316; **Fig. 5D**, lower panel).

SG-Transformer attention maps derived for each candidate among expert-labeled IEDs and non-IEDs (**Fig. 5E**, top panel) and probable ripples vs. probable IEDs (**Fig. 5E**, bottom panel). These conveyed which frequency band, at which times, contributed most to correct prediction. Generally, for IEDs and probable IEDs (**Fig. 5E**, right panels) the SG-Transformer model devoted attention to broadband frequencies as “triangular” spatial features spanning broad frequencies, remarkably consistent with that observed by eye in spectrogram representations of IEDs and other epileptiform activity in previous work^18,44^ and individual examples (**Fig. 1D**). Attention maps for non-IED ripple detections and probable ripples (**Fig. 5E**, upper and lower right panels) tended to emphasize spatial features circumscribed to the middle upper region of the spectrogram consistent with frequencies in the ripple band of range short duration, as would be expected based on previous work^2,3,6^ and visual inspection of individual examples (**Fig. 1C**).

Previous studies have demonstrated that ripples are predominantly generated in the CA1 region^3,45^ in both humans and rodents though from a detection standpoint they have been described in other subfields and even in the cortex.^3,46^ We thus evaluated the performance by focusing exclusively on IED/ripple candidates from the CA1 subfield of the hippocampus. Seven patients had imaging-confirmed electrode locations in the CA1 subfield, with 6,374 of the detected candidates encompassing this CA1 region. Limiting to only these candidates, we repeated the classification evaluations above, demonstrating negligible differences in results. Specifically, for expert-labeled data, the AUC-ROC score mean across five folds was 0.957 (range: 0.947–0.966), and PRAUC of 0.771 (range: 0.700–0.824) significantly above shuffled-label chance distributions (95% C.I.: 0.148∼0.152). For pseudo-labeled data, the AUC-ROC score achieved a mean of 0.918 (range: 0.889-0.934), and PRAUC of 0.916 (range: 0.905– 0.932) significantly above shuffled-label chance distributions (95% C.I.: 0.498-0.504).

### Leave-one-patient-out evaluations

We next aimed to evaluate the model’s generalizability to new (unseen) patients to understand its efficacy and translatability as a quantitative detector for research and clinical purposes. Using data from patients with expert-labeled hippocampal IEDs (n=15; **Fig. 4A**), we implemented the LOPO design (schematic in **Fig. 6A**), in which the model was trained on all but one of the patients and then tested on the held out (unseen) patient. We performed comparative analyses across three scenarios:

- Training and testing on expert labels (**Fig. 6B**)
- Training and testing on pseudo-labels (**Fig. 6C**)
- Training on pseudo-labels, and testing and testing on expert labels (**Fig. 6D**)

**Figure 6.**
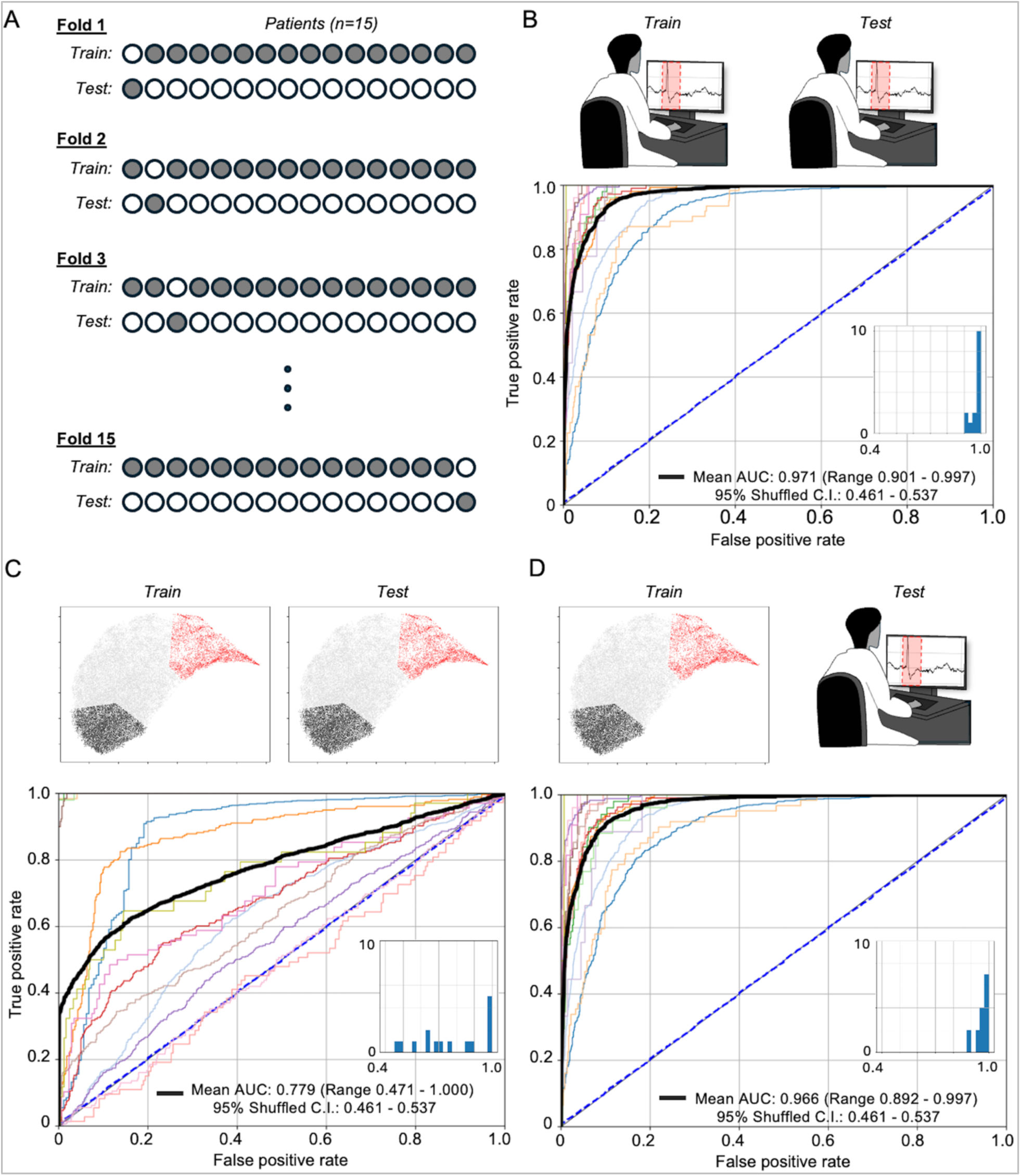
Leave-one-patient-out cross-validation SG-Transformer models. **A**. Schematic of the LOPO cross-validation method where, for each iteration, a model is freshly trained on data from 14 patients and tested on the remaining patient. **B**. AUC-ROC curves for training and testing on expert-labeled data (IEDs vs. non-IEDs), demonstrating excellent classification accuracy (mean AUC 0.971; inset: counts of individual patient AUCs). Lines correspond to individual patients (colors as in Fig. 1A). **C**. AUC-ROC curves for pseudo-labeled data, evaluated by training on the two poles of the UMAP distribution (Cluster 1 vs. Cluster 5, representing probable ripples and probable IEDs, respectively), demonstrating good classification accuracy (mean AUC 0.779). **D**. AUC-ROC curves for training on pseudo-labeled data (as in C), and testing on the expert-labeled data (as in B), again demonstrating excellent classification accuracy (mean AUC 0.966) comparable to if the model had been trained on expert-labeled data (as in B).

Training and testing on expert labels demonstrated an AUC-ROC mean across patient folds of 0.971 and range of 0.901-0.997 (**Fig. 6B**). The proportion of patients with an AUC significantly above chance based on 200 shuffled-label iterations was 100%. Training and testing on pseudo-labels demonstrated an AUC-ROC mean across patient folds of 0.779 and range of 0.471-1.0 (**Fig. 6C**). The proportion of patients with an AUC significantly above chance based on 200 shuffled-label iterations was 86.7%.

The third scenario of training on pseudo-labels and testing on expert labels, an evaluation of the degree to which classification by the SG-Transformer could replicate manual epileptologist-derived labeling of new (unseen) patients, was then performed. Training and testing on expert labels demonstrated an AUC-ROC mean across patient folds of 0.966 and range of 0.892-0.997 (**Fig. 6C**). The proportion of patients with an AUC significantly above chance based on 200 shuffled-label iterations was 100%. The probabilities derived for the individual candidates in this scenario were binned in UMAP space and illustrated in **Fig. 7A**, along with individual detection examples in **Fig. 7B**.

**Figure 7.**
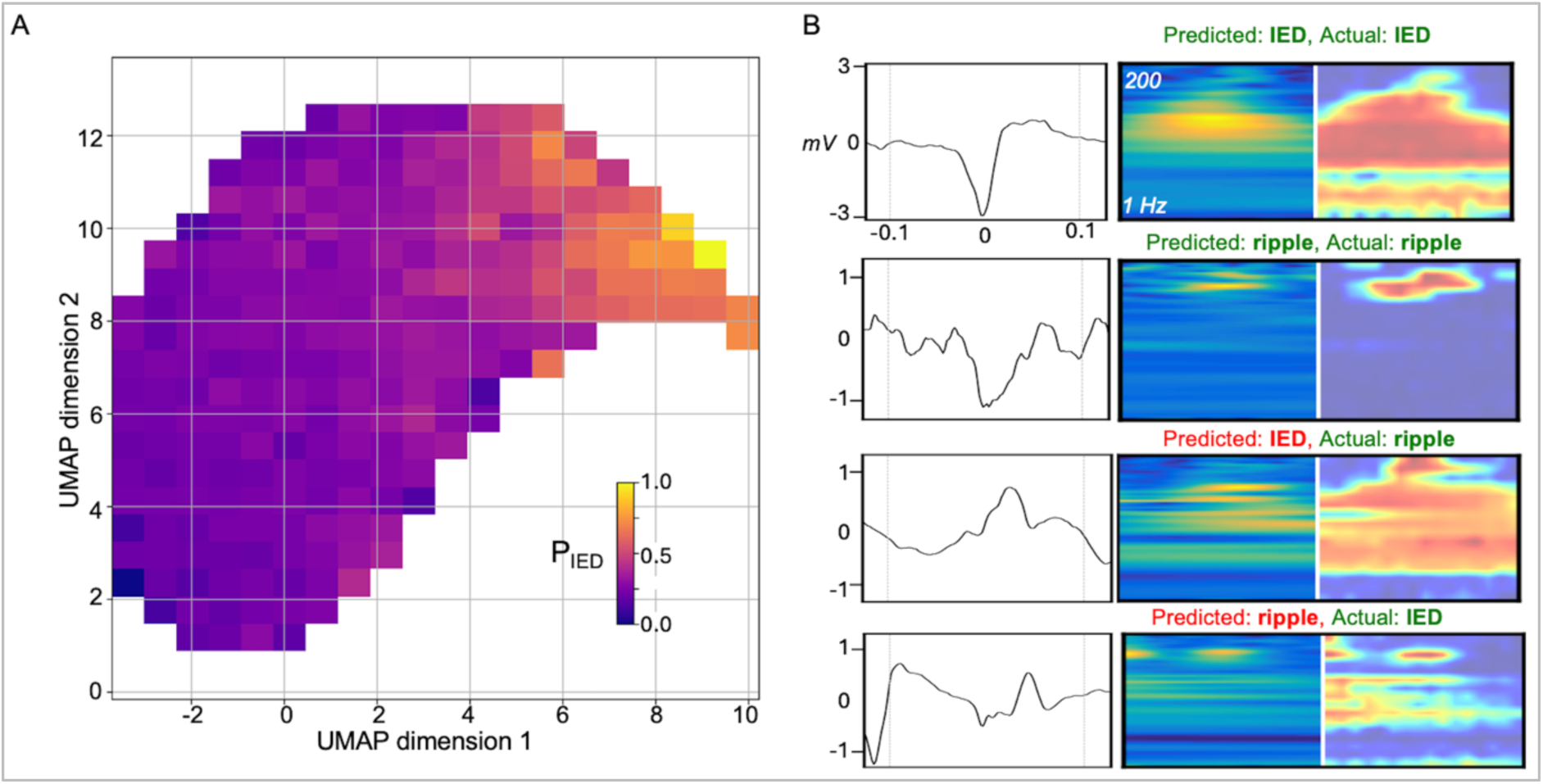
Gradient of IED probability and classification examples. **A.** IED probability (P_IED_, i.e. the likelihood of a candidate to be an IED, binned using mean) derived from LOPO cross-validation with training on pseudo-labels and testing on expert labels (data from Fig. 6D) illustrated as a gradient along original UMAP space. **B.** Representative examples (rows = individual detected candidates) of model performance showing candidate raw voltage traces at left, as well as their spectrograms and corresponding attention maps (middle and right panels; 200ms window, indicated by dashed line in raw traces at left). Examples from top to bottom are a true positive (IED), true negative (ripple/non-IED), false positive (likely related to substantial concurrent low frequencies as relayed by the attention map), and false negative (likely related to overlap of a potential ripple with the annotated duration of preceding sharp IED’s aftergoing slow wave).

## Discussion

We utilized unsupervised approaches (automated ripple detection and low-dimensional segmentation) to dissociate hippocampal ripples and IEDs along novel quantitative gradients,^33,47^ demonstrating that this approach may largely substitute the need for expert-labeled training data (**Fig. 6D**). Moreover, by implementing ViTs on spatial features of ICEEG spectrogram data, we achieved expert-level dissociation of ripples vs. IEDs.^37^ These combined approaches leverage the spatial (two-dimensional) and attention-based advantages of recent AI transformer architecture to extract the comprehensive spectrotemporal features that reliably distinguish normal from abnormal neural activity, rivaling human expert determinations. Importantly, we also demonstrate that the distinction of ripples and IEDs is best described as a continuum (i.e. a probability) rather than a binary categorization, on both group and individual-level data.

Detecting hippocampal ripples with standard filtering and thresholding approaches produces a massive parameter space of inherent spectral and temporal features across these “presumed ripple” candidates. Heterogeneity, such as different levels of power across lower (sub-ripple band) frequency ranges, drove the gradient space we observed for non-IED ripple candidates and suggests different but structured physiological implications across ripples, which may be key for understanding their physiological significance in human cognition. Moreover, epileptiform activity was easily drawn into the detections because these same filtering and thresholding techniques easily pick up on sharp activity through spectral leakage and other fleeting abnormal high-frequency activity events.^17,18^ Our unsupervised characterization of the massive assortment of over thirty thousand ripple candidates demonstrated no simple delineation between normal and abnormal, whether in raw, ripple-filtered, or spectrogram-based data substrates. We instead found a continuum, or a gradient space between the two, and this blurred overlap relates to the challenges of understanding the significance of these high frequency phenomena.

We used an unsupervised approach to understand these gradients inherent to hippocampal transient events by projecting this continuum of activity onto low-dimensional axes (UMAP). This framework enabled two advantages. First, pseudo-labels of presumed ripples and IEDs could be rapidly inferred,^47^ since epileptiform activity (verified by heavy overlap with expert-labeled events) was concentrated toward a tapered pole of the UMAP distribution. The k-means cluster that most clearly segmented these annotated events (Cluster 5), and its polar opposite (Cluster 1) manifested canonical epileptiform activity and physiological ripples, respectively. These labels indeed proved valuable in training and testing a classifier to distinguish between the two with moderate accuracy (**Fig. 5** and **Fig. 6C**).

Moreover, models trained on these pseudo-labels could distinguish expert-labeled IEDs from non-IEDs virtually as well as models trained on expert-labeled data (**Fig. 6D**). In other words, the model could predict human expert-quality labels across essentially all patients tested, and this finding was generalizable to new patients as assessed using a LOPO approach (**Fig. 6**). We attribute these performance results to the AI-based architecture of our SG-Transformer models focused on spatial features (relative time-frequency information) of comprehensive spectrogram representations. This comparable performance between the pseudo-label trained model and human/epileptologist annotations corroborates recent similar efforts used in scalp EEG^38–40,47–49^ and underscores emerging applications of quantitative spectrotemporal and other signal features in human intracranial neurophysiology.

While interpreting the key features in deep layers of existing AI models is challenging (e.g. CNNs), the attention-based vision-transformer approach allows extraction of spectrotemporal features (i.e. which frequences at which times) that were most important for classification. This avoids the problem of neural networks applied similarly, in which the relevant model weights are hidden and largely unobtainable in deep layers. In other words, the SG-Transformer enabled interpretable estimations of the time-frequency parameters that led to successful classification, an advantage that can be extremely difficult or impossible to replicate using CNNs and other AI classifiers.^50–52^

Our application of a classifier trained on canonical ripples and IEDs (Clusters 1 and 5) produced a quantitative estimation of the probability to which each candidate was estimated to be a spike (P_IED_, **Fig. 7A**). This represents continuous values as opposed to binary labels, and we speculate that they may enable a novel metric for future neurophysiological work. In other words, as opposed to most studies in which hippocampal ripples or IEDs are considered as distinct signal phenotypes, a quantitative metric like P_IED_ could be incorporated into statistical models to adjust for candidates that do not fit classic definitions yet may still have neurophysiological relevance.

To evaluate the spectrum of transient normal and abnormal events in human hippocampal recordings, we used emerging AI approaches to derive novel quantitative metrics. Rather than limit to classic features derived from raw voltage data (a 1-D vector representation), ViTs evaluated the comprehensive two-dimensional representations of neural activity encompassed by spectrograms. A primary advantage is the preservation of temporal (time) and spectral (frequency) contexts for classification, enabled by the SG-Transformer architecture. The 2-D spatial relations of spectral and temporal features provided a comprehensive quantitative backbone for our unsupervised analyses and classification results. Furthermore, these spectrotemporal-transforms of the data were key for stable representations. Namely, spectrograms were much more robust to small shifts in the temporal domain compared to using the raw or filtered waveforms (**Fig. 1E**). This simple factor is in fact crucial considering the simple shifts in alignment that can occur when deciding on re-centering procedures for detected events (e.g. ripples, IEDs, ERPs, etc.) and dramatically affect individual sample-by-sample features fed into quantitative models. A key reason underlying this advantage is that power or analytic amplitude spectrograms are phase-agnostic, reflecting frequency envelope changes which are less volatile than raw signals, thus reducing timepoint-by-timepoint variability that may undermine sample-by-sample precision requirements.

In addition, as opposed to CNNs which can also model two-dimensional data representations similarly yet with hidden layers, ViT models provide attention maps demonstrating the features of importance for classification results. Attention maps in this study clearly demonstrated the expected spectral leakage patterns for IED-like detections and the high gamma range “blobs” expected for physiological ripples.^18^

We used 61 logarithmically spaced frequencies from 1 to 200 Hz, over 103 time points, as the spectrogram data substrate of detected candidates. This preserved substantial spectral detail, with an intentional full broadband context over time for comprehensive low-dimensional projection, and for SG-Transformer based classification. Consolidating (e.g. averaging) these 61 frequencies into one or more canonical bands (e.g. alpha, gamma, etc.), as performed in most studies of hippocampal activity would simplify and improve computational speed but sacrifice detail and spectrotemporal context. We maintained detailed frequencies for two reasons. First, we suspected that preserving spectral detail would be crucial for ripple and IED classification. Indeed, UMAP clusters and SG-Transformer attention maps conveyed important components such as activity at distinct low frequencies that was important for unsupervised clustering (see non-IED clusters 1-4 in **Fig. 3G**) and classification based on attention maps (**Fig. 5E**). Second, canonical bands are generally based on scalp EEG, whereas intracranial signals including hippocampal oscillations have their own distributions of relevant frequencies. These are only partly defined in human work^53^ and chosen cut-offs even vary from study to study.^10^ Preserving spectral detail allowed us to remain agnostic to frequencies and derive the features that precipitated both in UMAP and the ViT models (attention maps), i.e. conveying which specific frequencies were important and at which times.

Our work has limitations. We kept our investigation focused on electrodes in the hippocampus given its well-established role in physiological ripples^54^ and its frequent implication in focal epilepsy.^7^ However, we anticipate that our findings may likely be generalizable to cortical ripples and IEDs given similar spectrotemporal profiles of both in cortical sites.^18,21,46^ We did not evaluate pathological events at higher frequencies such as fast ripples and HFOs.^19,21^ Such pathological phenomena will likely show distinct signatures when with spatial analysis of spectrotemporal data similar to the SG-Transformer approach applied herein, therefore related future investigations of epileptiform activity should include these higher frequency ranges. Review of examples of spectrogram attention maps **(Fig. 7B)** showed the SG-Transformer allocated attention to expected features based on a manual review, most notably triangular features of the spectrograms attributable to sharp edge artifact in IEDs.^6,18^ Yet important limitation is that such features would be specific to preprocessing approaches, as the use of the Hilbert or wavelet transforms preserves higher temporal detail at higher frequencies (hence triangular appearance to edge-related artifact) whereas the fast Fourier transform would entail a more vertical “pillar-like” artifact,^18^ thus we anticipate the signal processing tools and steps used for training and testing must be consistent.

Our work demonstrates that the contrast between ripples and IEDs is perhaps best described by a continuous gradient, due to the massive variety of sharpened waveform edges and pathological oscillations embedded with variable features of normal ripples that undermine ripple- or IED-focused investigations using typical detectors. Probabilities (i.e. of being a ripple) for each detection candidate could instead serve as a quantitative surrogate for ripples (i.e. 0 to 1, not 0 or 1) as a new framework to address and embrace the predominant gray zone between normal and abnormal high-frequency detections (**Fig. 3G**). This work also sets the stage for quantitative approaches leveraging ViT-based analysis of spatial information in intracranial spectrotemporal data, including much-needed interpretability of key features previously obscured in deep layers of traditional AI-based models.^42,47,48^ We anticipate these approaches will open new doors in future cognitive neurophysiology research and the detection of epileptiform activity in closed-loop or adaptive neuromodulation applications.

## Data availability

The data that support the findings of this study are available from the corresponding author, upon reasonable request.

## Acknowledgements

We thank Edward Chang and his laboratory for assistance with data collection. We also thank Yitzhak Norman, Daniel Alves Paladim, and Lip-Bu Tan for various technical input and support.

## Supplementary material

N/A

